# Measuring functional connectivity with wearable MEG

**DOI:** 10.1101/2020.09.25.313502

**Authors:** Elena Boto, Ryan M. Hill, Molly Rea, Niall Holmes, Zelekha A. Seedat, James Leggett, Vishal Shah, James Osborne, Richard Bowtell, Matthew J. Brookes

**Affiliations:** Sir Peter Mansfield Imaging Centre, School of Physics and Astronomy, University of Nottingham, University Park, Nottingham, NG7 2RD, United Kingdom; QuSpin Inc. 331 South 104^th^ Street, Suite 130, Louisville, 80027, Colorado, USA

**Keywords:** Optically-pumped magnetometer, OPM, Magnetoencephalography, MEG, OPM-MEG, functional connectivity, network, wearable MEG, amplitude-envelope correlation, AEC

## Abstract

Optically-pumped magnetometers (OPMs) offer the potential for a step change in magnetoencephalography (MEG) enabling wearable systems that: provide improved data quality; accommodate any subject group; allow data capture during movement and offer a reduction in costs. However, OPM-MEG is still a nascent technology and, to realise its potential, it must be shown to facilitate key neuroscientific measurements, such as the characterisation of human brain networks. Networks, and the connectivities that underlie them, have become a core area of neuroscientific investigation, and their importance is underscored by many demonstrations of their perturbation in brain disorders. Consequently, a demonstration of network measurements via OPM-MEG would be a significant step forward. Here, we aimed to show that a wearable 50-channel OPM-MEG system enables characterisation of the electrophysiological connectome. To this end, we characterise connectivity in the resting state and during a simple visuo-motor task, using both OPM-MEG and a state-of-the-art 275-channel cryogenic MEG device. Our results show that connectome matrices from OPM and cryogenic systems exhibit an extremely high degree of similarity, with correlation values >70 %. This value is not measurably different to the correlation observed between connectomes measured in different subject groups, on a single scanner. In addition, similar differences in connectivity between individuals (scanned multiple times) were observed in cryogenic and OPM-MEG data, again demonstrating the fidelity of OPM-MEG data. This demonstration shows that a nascent OPM-MEG system offers results similar to a cryogenic device, even despite having ∼5 times fewer sensors. This adds weight to the argument that OPMs will ultimately supersede cryogenic sensors for MEG measurement.

## 1 Introduction

Since its inception, functional neuroimaging has made many important contributions to our understanding of brain function, and one of the most significant is the discovery of brain networks. A network is found when a statistical relationship between neuroimaging signals, derived from spatially separate brain regions, is shown to exist. Such a relationship is termed functional connectivity. The first measurements of functional connectivity used functional magnetic resonance imaging (fMRI; Biswal et al., (1995)) to measure correlation between blood-oxygenated-level-dependent (BOLD) time courses from left and right motor cortex, in the absence of a task (in the so-called resting state). Following this, many fMRI studies (e.g. (Beckmann et al., 2005; Fox and Raichle, 2007; Smith et al., 2009)) focused on identifying other resting-state networks (RSNs); some associated with sensory processing (e.g. auditory or visual networks) and others with attention and cognition (e.g. the default-mode and dorsal-attention networks). Raichle, (2009) described this era of functional imaging as a "paradigm shift". Indeed, study of RSNs offers a powerful means to investigate healthy function, and dysfunction in a wide range of disorders, including schizophrenia, depression, anxiety and dementia (Menon, 2011).

Most functional connectivity studies have been based on fMRI. However, the BOLD signal is an indirect metric of function, based on haemodynamics, which leads to significant disadvantages: for example, Bright and colleagues (Bright et al., 2020) showed an overlap between neural and vascular network components and this makes the interpretation of fMRI networks challenging without first understanding network-specific vascular architecture. In addition, whilst fMRI exhibits exquisite spatial resolution (∼1 mm accuracy), it has extremely limited temporal resolution due to the latency and longevity of the haemodynamic response. This means that the timescale of a functional connectivity measurement is limited, and so it is challenging to probe the formation and dissolution of functional networks on a time scale relevant to cognition (Hutchison et al., 2013). For these reasons, a move towards electrophysiological imaging techniques (which exhibit significantly better temporal resolution) for the characterisation of network connectivity is extremely important.

Magnetoencephalography (MEG; Cohen, (1972)) measures the magnetic fields generated outside the head by current flow through neuronal assemblies in the brain. In this way, it offers a means to bypass haemodynamics and infer directly the electrophysiological connectome. MEG data (like all electrophysiological data) are dominated by “neural oscillations” (synchronised rhythmic electrical activity across neurons) in the 1–200 Hz frequency range. Emerging evidence suggests that these oscillations mediate (at least in part) network formation and consequently their measurement offers an exciting means to probe intrinsic network coupling (Engel et al., 2013). MEG has been successfully used to measure functional connectivity in many studies (e.g. (Baker et al., 2014; Brookes et al., 2012, 2011; Gross et al., 2001; Hipp et al., 2012; Luckhoo et al., 2012; O’Neill et al., 2015)). Networks similar to those seen with fMRI have been and the excellent temporal resolution that MEG affords enables measurement of dynamic connectivity. Indeed, recent studies show that canonical networks modulate on the time scale of seconds (O’Neill et al., 2015) and even milliseconds (Baker et al., 2014). More recently, Seedat et al., (2020) suggested the involvement of extremely short (e.g. ∼300 ms) punctate events in driving canonical network connectivity, further underscoring the importance of temporal precision in the characterisation of the human connectome.

A combination of high spatial and temporal accuracy means that MEG offers arguably the best means to measure functional connectivity and the networks that it underpins. However, existing MEG systems have huge limitations: the sensors that form the basic building block of MEG systems (superconducting quantum interference devices; SQUIDs) operate at cryogenic temperatures. These sensors must therefore be fixed in position within a cryogenic dewar, making systems large, cumbersome, and “one-size-fits-all” – i.e. they cannot adapt to different head shapes or sizes. This results in inhomogeneous and sometimes poor brain coverage (particularly in infants). Even if the head is well fitted to a MEG system, the gap between the scalp and the sensors that is needed for thermal insulation reduces sensitivity. The fixed nature of the sensors means that any head movement during data acquisition significantly reduces data quality. Subjects must therefore remain extremely still, which makes the environment poorly tolerated by many groups, including children. Finally, the cryogenic infrastructure and complex electronics makes systems expensive. These factors have, to date, limited the uptake of MEG as a neuroimaging modality, and if MEG-based connectome measures are to realise their potential for neuroscientific discovery and clinical translation, then new types of MEG technology will be required.

Recent advances in quantum technology have led to the development of a new type of magnetic field sensor. Optically-pumped magnetometers (OPMs) offer measurement of magnetic field with a similar sensitivity to the cryogenic sensors used in conventional MEG, however they do not require cooling. Furthermore, they are very small, and lightweight. This has led a number of groups to begin to fabricate OPM-based MEG devices. Suitability of OPMs to capture neuromagnetic signals has been well documented (e.g. (Barry et al., 2019; Borna et al., 2020; Boto et al., 2017; Iivanainen et al., 2019; Johnson et al., 2013; Kamada et al., 2015; Kim et al., 2014; Roberts et al., 2019; Sander et al., 2012; Tierney et al., 2018; Xia et al., 2006) and more recently their lightweight nature has been exploited to develop “wearable” systems in which (if background fields are controlled appropriately) subjects can move their head freely during data acquisition (Boto et al., 2018). OPM arrays are beginning to be developed with up to 50 sensors surrounding the head (Hill et al., 2020) and there is a growing argument that these devices – which are also cheaper than conventional MEG – will ultimately supersede the current generation of MEG scanners. However, OPM-MEG remains a nascent technology and if the functional neuroimaging field is to gain confidence in it, OPM-based systems must be able to do everything a SQUID system can do. Given the importance of functional connectivity, demonstration of its measurement with OPM systems is a vital step.

There are a number of reasons why functional connectivity is a challenge for OPM-MEG. Firstly, most OPM experimental demonstrations have targeted specific brain regions due to a relatively low sensor count. Given that networks are distributed across the whole brain, coverage that rivals conventional MEG is required. Secondly, networks are highly spatially-specific and their measurement consequently relies on good spatial resolution; if an OPM system is to characterise these networks then it must also offer similar or better spatial specificity to conventional MEG scanners. (At present, this would mean achieving similar spatial resolution with around 50, rather than ∼300 sensors). Finally, functional connectivity is heavily reliant on high quality data since, unlike task-based studies where data can be averaged over many trials (and thus artefacts will often be averaged out) functional connectivity must be measured using unaveraged data. This latter point is amplified since unlike conventional MEG systems which often rely on a gradiometer formulation to reduce environmental interference, OPMs are naturally formed as magnetometers. This, at least in principle, increases the effect of external interference, both from environmental (e.g. lab equipment) and biological sources (e.g. the heart).

In this paper, we aim to test whether a 50-channel OPM-MEG system can successfully measure the functional connectome. We measure connectivity during a simple motor task, and in the resting state, and in both cases we compare our findings to those generated using a 275-channel cryogenic system (which for the purposes of this study will be treated as a “gold standard”).

## 2 Methods

### 2.1 OPM-MEG system

The wearable OPM-MEG device used in this study has been developed at the Sir Peter Mansfield Imaging centre, University of Nottingham, and was described recently in Hill et al., (2020). A schematic of the system is shown in Figure 1a: the OPM-MEG suite contains a magnetically-shielded room (MSR), who’s design has been optimised for OPM operation (MuRoom, Magnetic Shields Ltd. Kent, UK). This MSR provides a remnant magnetic field magnitude <2 nT and <2 nT/m magnetic field gradient following a demagnetisation procedure (Altarev et al., 2015). These fields are significantly lower than those found in MSRs which do not feature demagnetisation coils.

**Figure 1:**
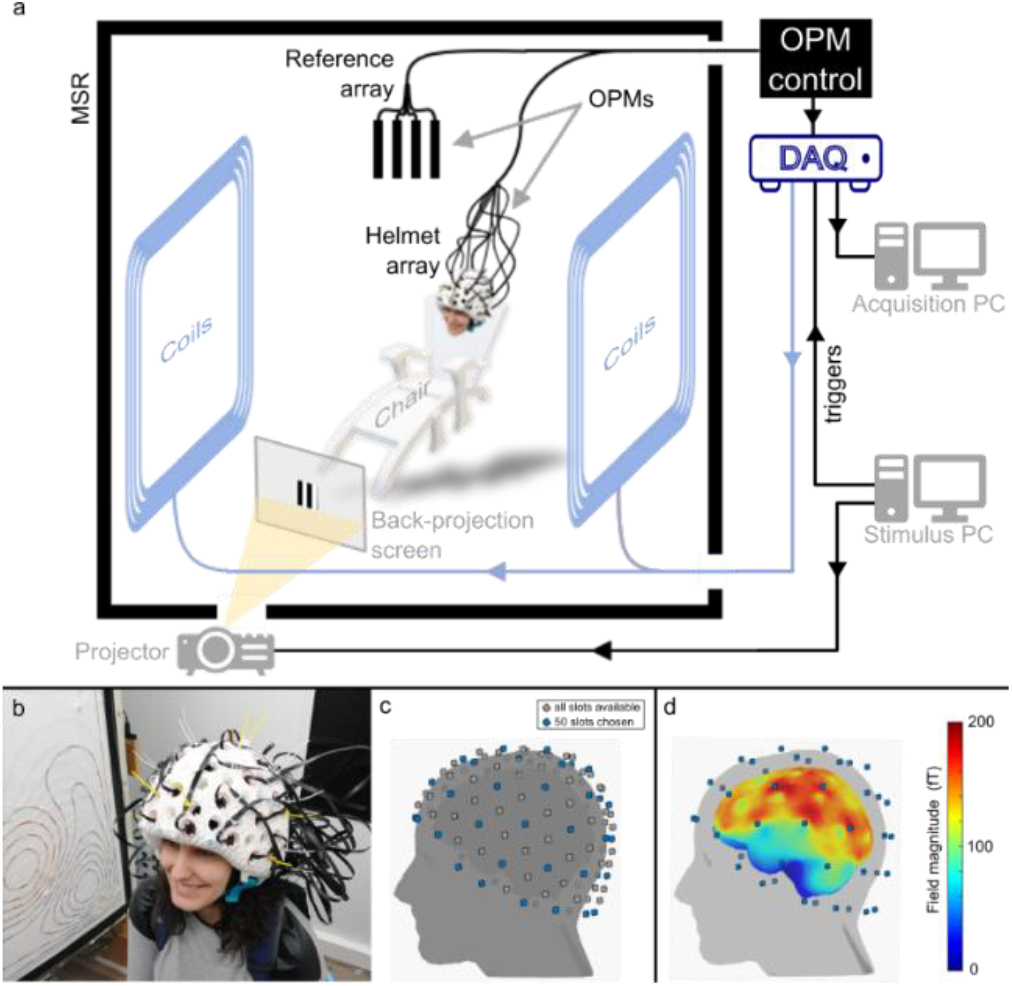
The OPM-MEG system. a) Schematic of the OPM-MEG suite. b) Photograph of subject wearing an additively-manufactured helmet with 50 OPM sensors mounted within it. c) Digitised head surface for an example participant, showing the 133 slots available in the helmet (grey) and the 50 chosen for this study (blue). Note that OPMs were made sensitive to the field in the radial direction only. d) Cortical coverage achieved by the selected 50 OPM locations: the norm of the forward fields across all sensors is plotted at each vertex of the brain surface.

Inside the MSR, the participant sits in a non-magnetic chair, and wears an additively manufactured rigid helmet in which 50 OPMs (Gen-2 zero-field magnetometers, QuSpin Inc., (Osborne et al., 2018)) are mounted (see photograph in Figure 1b). The helmet contains a total of 133 possible slots where OPMs can be placed (grey dots in Figure 1c), and for the study described in this manuscript, we chose 50 locations to provide whole-head coverage (blue dots). The OPMs themselves were configured to measure the radial component of magnetic field. Figure 1d shows the cortical coverage achieved with 50 radial OPMs placed in the slots shown in blue: for each vertex of the brain surface, the colour represents the norm of the field (across all sensors) produced by a current dipole located at that vertex. (Dipoles were oriented in two orthogonal tangential directions, and the result is taken to be the average of the two lead field norms.) Note that reasonable coverage of the whole cortex is achievable, although sensitivity to the temporal pole is somewhat diminished (a problem also found in conventional MEG devices).

Further reduction of background fields is achieved with a set of bi-planar coils (Holmes et al., 2018) positioned at either side of the participant, coupled to an OPM reference array placed behind the subject’s head. The remnant static magnetic field and its spatial variation inside the MSR is estimated by the reference array, and an equal and opposite field applied by the coils in order to effect field cancellation. This reduces the remnant field, better enabling OPM operation, minimising any artefact caused by the subject moving their head (and consequently the sensors) through the background field.

The OPMs themselves have been described previously and a complete description will not be repeated here (see e.g. Tierney et al., (2019) or Boto et al., (2020) for a review). Note that they are controlled via a computer located outside the MSR, and analogue output signals directly proportional to local magnetic field, are fed into a National Instruments (Austin, TX, USA) Digital Acquisition unit (DAQ), digitised, and recorded. A separate computer is coupled to stimuli delivery systems, and sends triggers to the DAQ which are synchronously recorded with the MEG data.

### 2.2 OPM Co-registration

In order to enable source localisation, accurate knowledge of the sensor locations and orientations relative to the brain is required. This was provided by a 3-dimensional optical imaging system (structure IO camera (Occipital Inc., San Francisco, CA, USA) coupled to an Apple iPad, operating with Skanect Pro software) and an anatomical MRI scan (Hill et al., 2020; Homölle and Oostenveld, 2019; Zetter et al., 2019). The anatomical MRI was recorded using a 3 T Philips Ingenia MRI system, running an MPRAGE sequence, at an isotropic spatial resolution of 1 mm. The locations and orientations of the sensor casings with respect to the helmet are known *a-priori* from the additive manufacturing process, and the co-registration procedure was used to map the helmet on to the head. This was done in two stages. First, 6 coloured markers were placed at known locations on the helmet, with a further 4 on the participant’s face. The camera, coupled with a colour-thresholding algorithm, was used to map the relative locations of these markers, allowing mapping of the helmet to the face. Following this, the helmet was removed and the participant was asked to wear a swimming cap (to flatten their hair). A second digitisation was then acquired measuring the positions of the markers on the face, relative to the rest of the head surface. The head surface was then fitted to the equivalent surface extracted from the anatomical MRI scan. Combining two transforms (helmet-to-head and head-to-MRI) we were able to effect a complete co-registration of sensor casing to brain anatomy. The location of the sensitive cell within the OPM casing was accounted for and we assumed that the sensitive axis was radial, and parallel to the external sensor housing.

### 2.3 Task-based connectivity experiment

#### 2.3.1 Paradigm and data acquisition

Two subjects undertook a visuo-motor task. The task comprised presentation of a centrally-presented, inward-moving, maximum-contrast circular grating (Hoogenboom et al., 2006; Iivanainen et al., 2019), which is known to increase gamma oscillations in the visual cortex. Whilst the visual stimulus was on the screen, participants were asked to perform continuous abductions of their right index finger; a task known to modulate beta oscillations in sensorimotor cortex. The grating was presented for either 1.6, 1.7 or 1.9 s. Each trial ended with a 3-s baseline period, and 100 trials were recorded.

Both participants were scanned six times in the OPM-MEG system and six times in a cryogenic MEG instrument (CTF, Coquitlam, BC, Canada). The study was approved by the University of Nottingham Medical School Research Ethics committee.

OPM data were acquired using a sampling frequency of 1,200 Hz. 42 and 49 OPM sensors were available for participants 1 and 2, respectively. The visual stimulus was back projected onto a screen placed ∼85 cm in front the participant’s head. A separate co-registration procedure was performed after each experiment.

Cryogenic MEG data were acquired using a 275-channel CTF system, operating in 3^rd^-order gradiometer configuration, at a sampling frequency of 600 Hz. The stimulus was presented on a back-projection screen placed 95 cm in front of the participant (note that the stimulus was matched for visual angle between the two scanner types). Three head-position indicator (HPI) coils were placed on the participant’s head at three fiducial locations (nasion, left and right pre-auricular points). Continuous tracking of the head was achieved via these coils, which were periodically energised during acquisition. A 3D digitiser (Polhemus) was used to measure the locations of the fiducial markers relative to the head surface, prior to each experiment. By matching the participant’s digitised head surface with the equivalent surface extracted from their anatomical MRI, co-registration of the fiducial markers, and consequently the MEG sensor geometry, to the individual’s brain anatomy was achieved.

#### 2.3.2 Data analysis

All MEG data (OPM and cryogenic) were bandpass filtered between [8–80] Hz and epoched into trials. “Bad” trials were removed in cases where the standard deviation of the signal was greater than 3 times the average standard deviation across all trials. This procedure was carried out independently for each channel. Data were then concatenated into a single signal per channel.

Both cryogenic and OPM data were analysed in the same way: functional connectivity was calculated between 78 discrete cortical regions, defined based on the automated anatomical labelling (AAL) atlas (Tzourio-Mazoyer et al., 2002). A scalar beamformer was used to obtain a single electrophysiological time course representative of each region (i.e. a ‘virtual electrode’ placed at the centre of mass of each region). The data covariance matrix was computed in the 8–80 Hz frequency range for a time window spanning the whole experiment. Regularisation was not applied, to maximise spatial resolution (Brookes et al., 2008). The forward model was based on a single sphere for the OPM system and multiple local spheres for the cryogenic system.

After beamforming, regional signals were frequency-filtered to the alpha (8–13 Hz), beta (13–30 Hz), and gamma (52–80 Hz) frequency bands, and epoched into trials. Pairwise orthogonalization (Brookes et al., 2012; Hipp et al., 2012) was used to mitigate the problem of signal leakage between AAL regions (itself a result of the ill-posed MEG inverse problem). Following this, the absolute value of the Hilbert transform of the frequency filtered data was computed to generate the amplitude envelope of oscillatory signals, which was then down-sampled to 10 Hz. Pearson correlation was calculated between amplitude signals for all possible AAL region pairs and averaged over trials. For each participant, each experiment and each frequency band, this procedure resulted in a single 78 x 78 adjacency matrix (or connectome) representing whole-brain connectivity. Finally, connectivity matrices were averaged across experimental runs.

OPM and cryogenic results in the beta band were compared. (Note we chose the beta band as this range has been shown to provide robust brain network measurements (e.g. (Hunt et al., 2016)). Comparisons were made in two ways:

- **Within- and across-subject correlation:** To quantitatively compare connectivity matrices between MEG systems, we first vectorised the matrices from all experimental runs. For each subject, the 6 OPM runs and 6 cryogenic runs were paired (randomly) and Pearson correlation between the vectorised adjacency matrices was calculated, and averaged. In this way, we obtained a within-subject correlation between OPM-derived and cryogenic-derived connectivity matrices. We also performed the same calculation between subjects (e.g. correlating connectivity from subject 1’s cryogenic data and subject 2’s OPM data). We hypothesised that there would be individual differences in the connectivity matrices between subject 1 and subject 2, and that these differences would be maintained across the two scanner types (i.e. colloquially, ‘the scanner would know who it was scanning’). Consequently, we hypothesised that correlation values would be higher within subject than between subject. A Wilcoxon sign rank test was performed to assess statistical significance.
- **Connectivity strength:** We calculated the linear sum of elements within each connectivity matrix, in one direction; this resulted in 78 regional values of connectivity strength (i.e. for each of the AAL regions, this metric represents the strength of the connection between that region and all other regions in the AAL atlas). Connectivity strength was separately calculated for each subject, MEG system, and experimental run. Again, we wished to probe whether individual differences in connectivity strength were maintained across the two MEG systems. To this end, for each of the 78 regions, a t-test was used to determine the statistical significance of differences between subjects. These calculations were performed for each scanner type separately and multiple comparisons (across the 78 regions) were controlled using the Benjamini-Hochberg procedure (Benjamini and Hochberg, 1995). We hypothesised that any regions where a significant between-subject difference occurred, would be matched across scanner types.

### 2.4 Resting-state connectivity experiment

#### 2.4.1 Paradigm and data acquisition

Seven subjects (2 females, mean age 25.86 ± 4.30) took part in the resting-state study. All participants gave written informed consent, and the study was approved by the University of Nottingham Medical School Research Ethics Committee.

Seven minutes of eyes-open, resting-state MEG data were acquired using the wearable OPM-MEG system at a sampling frequency of 1,200 Hz. Participants were asked to fixate on a small red cross which was centrally positioned on a grey background on the back-projection screen. Apart from this, they were simply asked to relax and do nothing. Participants were free to move during the recording, but they were not encouraged to do so. Co-registration was performed (as described above) at the end of each experiment, and MRI’s were available for all participants.

For comparison, we employed eyes-open resting-state data which had been acquired previously, in 63 subjects as part of the United Kingdom MEG Partnership (UKMP; (Hunt et al., 2016)) programme. These data were all acquired using the same 275-channel cryogenic MEG system used in our task-based study. The system was operated in third-order synthetic gradiometer configuration, and data were acquired at a sampling frequency of 1,200 Hz. The paradigm comprised a 5-min recording during which participants focused on a small, centrally-positioned red circle, back-projected onto a screen. Head position monitoring was facilitated via three head-position indicator coils which were energised during the scan. Co-registration of MEG sensor geometry to individual brain anatomy was achieved via head shape digitisation.

#### 2.4.2 Data analysis

Data were band-pass filtered between 8 and 80 Hz, and bad channels were discarded based on visual inspection. This meant there were OPM data from 49, 47, 47, 48, 47, 48 and 45 channels for subjects 1–7, respectively.

OPM and cryogenic data were processed in the same way. A scalar beamformer was employed to reconstruct a representative signal at the centre of mass of each of the 78 AAL regions. Data covariance was computed in the 8–80 Hz frequency band and within a time window encompassing the complete resting state recording. Regularisation was not used (see also appendix). Following this, regional signals were frequency-filtered into the alpha (8–13 Hz) and beta (13–30 Hz) bands, and pairwise orthogonalisation used to mitigate signal leakage. The absolute value of the analytic signal was computed for each regional time course, to generate the amplitude envelope of oscillatory signals, which was then down-sampled to 5 Hz. Pearson correlation was calculated between envelopes for each region pair. For each participant and frequency band, this resulted in a single connectivity matrix representing whole-brain connectivity. Group connectivity matrices were computed by averaging across subjects.

We aimed to show that OPM derived connectivity was similar to connectivity derived using a cryogenic instrument and to this end we exploited the large UKMP dataset. We randomly grouped the cryogenic derived connectivity matrices from our 63 subjects into 9 groups, with 7 subjects per group. For each frequency band, we computed a group average connectivity matrix. This resulted in nine matrices derived from cryogenic data which we could compare with the single average (also of 7 subjects) derived from our OPM system.

Quantitative comparison was made by correlating the vectorised connectivity matrices. For each frequency band, we first derived correlations between the OPM group average, and each of the nine cryogenic matrices. For comparison, we then measured correlation, in the same way between all possible pairs of cryogenic-derived connectivity matrices (i.e. group 1 to group 2; group 1 to group 3 etc.). This yields a total of 36 correlation values showing how different, randomly selected, groups compare in terms of their whole brain connectome. We reasoned that if OPM-derived connectivity was significantly different to cryogenic-derived connectivity then we would expect OPM-to-cryogenic correlation to be lower than cryogenic-to-cryogenic. Obviously these 36 values are not statistically independent quantities. For this reason, to test this statistically, we randomly selected 9 independent comparisons from our 36 group pairings, and, via a Wilcoxon sign rank test, we tested for a significant difference between these 9 values (which represent cryogenic-cryogenic group similarity), and the equivalent 9 OPM-cryogenic correlation values.

## 3 Results

### 3.1 Task-based connectivity

Figure 2 shows results from our task-based connectivity study. For each participant, connectivity matrices obtained from OPM and cryogenic data averaged across six experimental runs, are shown. The left panel shows results in the alpha band, the middle panel shows beta band and the right panel shows gamma band. Colour indicates connectivity values. The inset 3D brain plots show the dominant connections between AAL regions (represented by the red lines) that lie within 35 %, 45 % and 45 % of the maximum connectivity value in alpha, beta and gamma, respectively. Clear differences in network structure can be seen between the three bands: the alpha-connectome is predominantly occipital, although subject 1 shows some parietal connections; the beta band shows primarily parietal (bilateral sensorimotor) connections. Anecdotally, we note a more unilateral network in subject 1 and a bilateral network in subject 2. Finally, the gamma-band connectome is dominated by occipital connections. Similarities between OPM- and cryogenic-derived matrices are clear, a good example being the agreement on inter-individual differences that are shown in the beta band.

**Figure 2:**
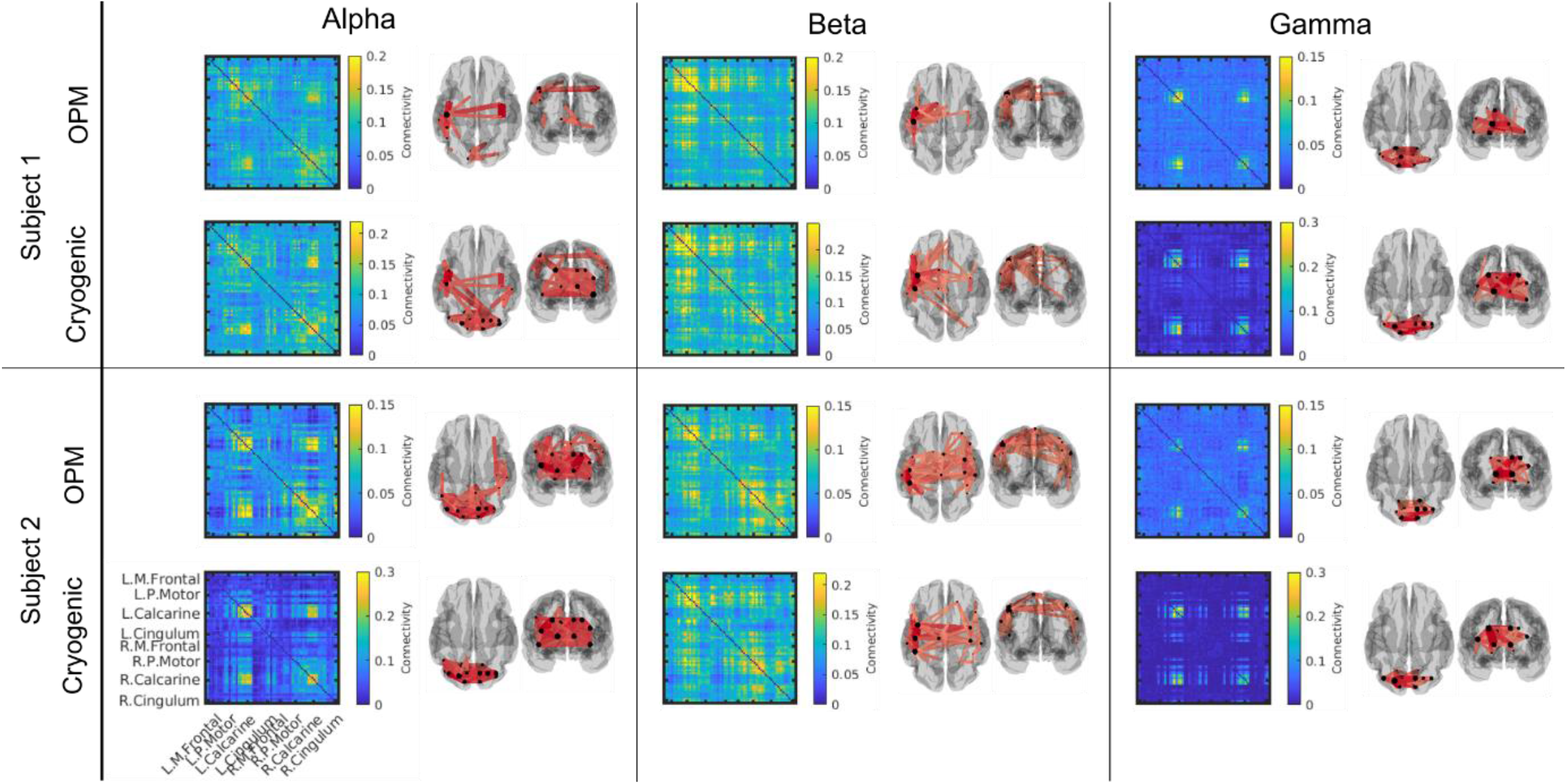
Task-based functional connectivity matrices. Average connectivity matrices (across 6 runs) in the alpha (left), beta (middle) an d gamma (right) bands for participants 1 (top) and 2 (bottom). For each participant, both OPM-derived (top) and cryogenic-derived (bottom) matrices are shown. Colour bars show Pearson correlation values. Alongside the matrices, the 3D brains show dominant connections (within 35 % (alpha), 45 % (beta) and 45 % (gamma) of the maximum value).

Figure 3 probes the similarity of connectome matrices, within- and between-subjects, in OPM and cryogenic recordings. Comparison is made in the beta band only. In panel a, scatter plots show OPM-derived connectivity values for all region pairs, plotted against the equivalent cryogenic-derived connectivity values i.e. each point on the graph represents a measured connection, and assuming OPMs and cryogenic sensors measure the same connectivities, in the same subject, we would expect to see a linear relationship. Plots on the left show a within-subject comparison: the top scatter plot, in blue, corresponds to subject 1 and the bottom plot, in yellow, corresponds to subject 2. The scatter plots on the right compare connectivity matrices between subjects. Note that to generate these plots, we have randomly paired and overlaid OPM and cryogenic runs. (e.g. OPM run 1 plotted against cryogenic run 3; OPM run 2 plotted cryogenic run 5, and so on; 6 independent comparisons are overlaid). A line of best fit is also added and the dotted line shows ‘*y* = *x*’.

**Figure 3:**
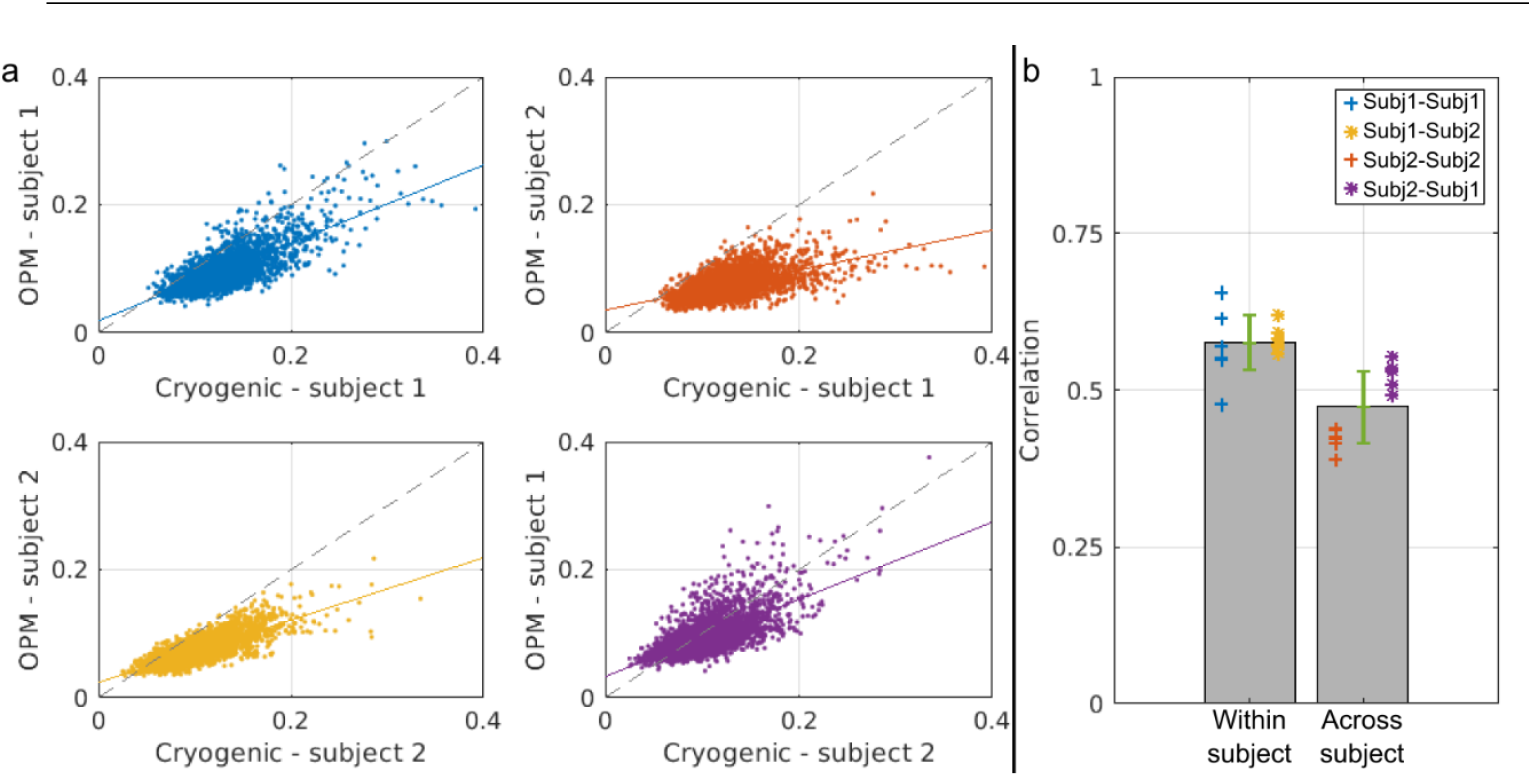
Cryogenic vs OPM connectivity in the beta band. a) Scatter plots showing connectivity values derived from cryogenic data plotted against connectivity values derived from OPM data (each ‘dot’ depicts a measured connection). Left column shows within-subject correlation for subject 1 (top) and subject 2 (bottom). Right column corresponds to between-subject correlation: b) Bar plot showing the mean within- and between-subject correlation values. Error bar corresponds to standard deviation across the 6 comparisons. Crosses and stars indicate individual values.

Note first that a clear linear relationship is observed, demonstrating that OPM- and cryogenic-derived connectivity matrices are similar in structure. Note also that within-subject correlation (left-hand scatter plots) appears tighter than between-subject correlation (right-hand scatter plots). The bar chart in Figure 3b directly illustrates this point. The bar chart shows mean within-subject correlation (0.58 ± 0.04) and between subject correlation (0.48 ± 0.07). Error bars correspond to standard deviation. The crosses and stars show the data that contribute to these averages. The blue crosses show correlation between OPM and cryogenic connectomes, within subject 1; the yellow stars show the same thing, for subject 2. Red crosses show the subject 1 OPM connectome versus the subject 2 cryogenic connectome; purple stars show the subject 2 OPM connectome versus the subject 1 cryogenic connectome. Within-subject correlation was significantly (*p* = 0.031) higher than between-subject correlation, lending support to our hypothesis that individual differences between subjects have been replicated in measurements made using the two types of scanner.

Results of our connectivity strength analysis are shown in Figure 4. Panel (a) shows line plots depicting normalised connectivity strength from the cryogenic- (red) and OPM-derived (blue) beta-band connectome, for subject 1 (top) and 2 (bottom). On the x-axis each of the AAL regions are represented. Thick lines correspond to the average connectivity strength and shaded areas represent standard deviation, across all six runs. There are clear similarities between the cryogenic- and OPM-derived values: firstly, the regions with highest values correspond to sensorimotor areas (which is to be expected given the task). Secondly, both scanners show a clear difference between subjects around the right motor regions. This is better visualised in Figure 4b, where the normalised connectivity strength is plotted on a 3D brain. Cryogenic results are at the top, OPM results at the bottom, left panels correspond to subject 1 and right panels to subject 2. For each subject, both cryogenic and OPM data yield very similar connectivity strength patterns Again, subject 1 exhibits a more unilateral beta-band connectome, as opposed to subject 2, in which a clear bilateral network can be observed. Panel (c) shows the same data as in panel (a) but grouped by scanner type: OPM at the top, cryogenic at the bottom. In both plots, subject 1 is represented with a solid line and subject 2 with dashed line. Here, the difference in connectivity strength between both participants around the right sensorimotor areas can be seen clearly. Figure 4d shows brain regions whose connectivity strength differed significantly (*p* < 0.05 corrected) between subjects. Both cryogenic (bottom) and OPM (top) data highlight similar regions – in particular right sensorimotor cortices and premotor areas stretching forward to the frontal lobe.

**Figure 4:**
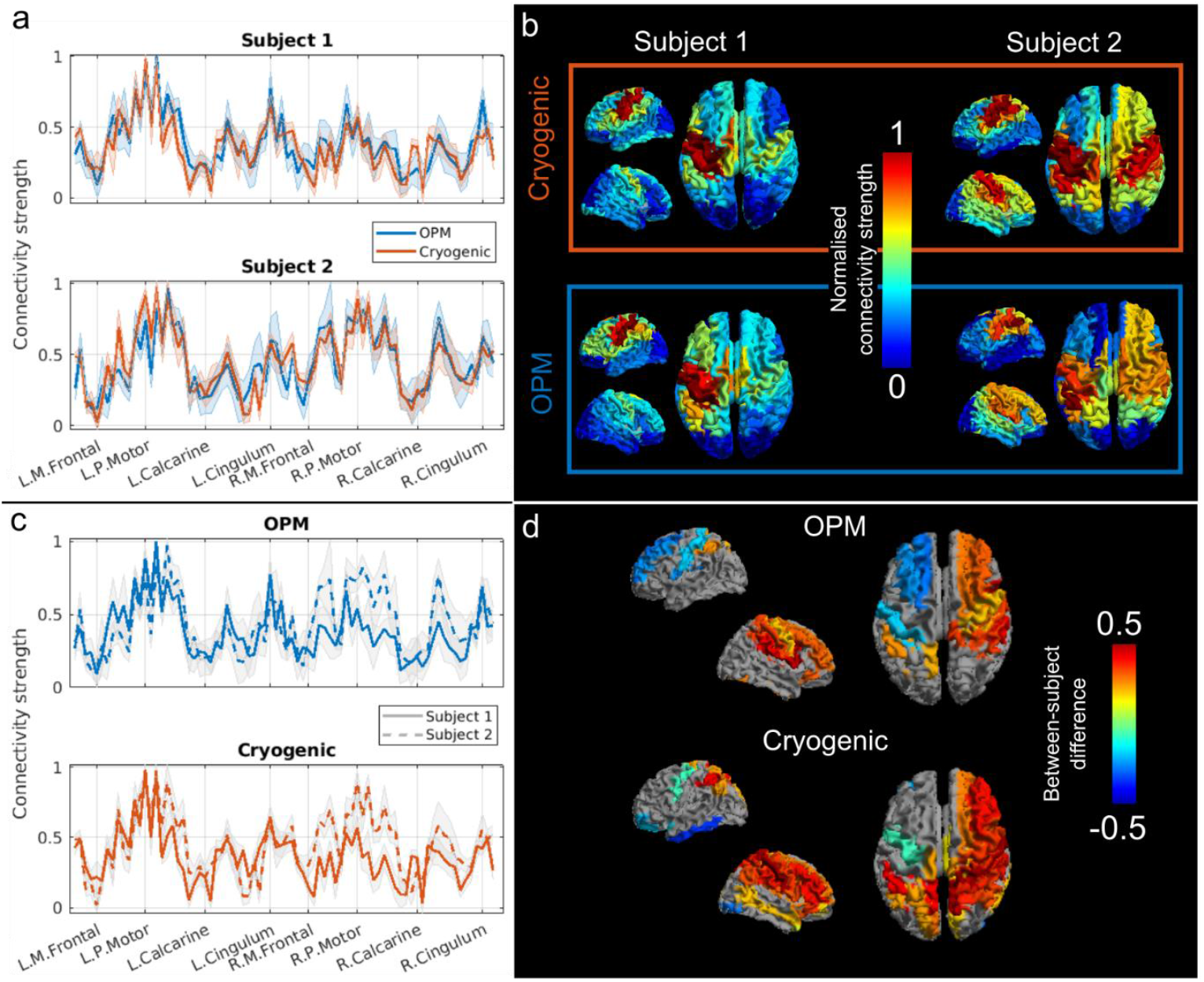
Connectivity strength in the beta band. a) Normalised connectivity strength recorded using cryogenic- (red) and OPM- (blue) derived data. Values are plotted for all 78 AAL regions, for participants 1 (top) and 2 (bottom). The shaded area represents standard deviation across 6 runs. Note the similarities between cryogenic and OPM plots. b) Normalised connectivity strength plotted on the brain surface for both subjects and both systems. c) Same as (a) but grouped by scanner type: normalised connectivity strength recorded using cryogenic- (bottom) and OPM- (top) derived data for participants 1 (solid line) and 2 (dashed line). d) Brain areas showing significant difference between participants (grey indicates no significant difference). Note both systems highlight similar regions.

### 3.2 Resting-state connectivity

Results from the resting-state OPM-MEG connectivity study are shown in Figure 5. Group-average connectivity matrices in the alpha (panel a) and beta (panel b) bands are plotted. The 3D brain plots show dominant connections between AAL regions (connections which lie within 20 % of the maximum). Differences between alpha- and beta-band connectomes are clear; alpha oscillations support connections between occipital and motor regions (with some frontal projections), whilst the beta-band connectome appears dominated by a sensorimotor network.

**Figure 5:**
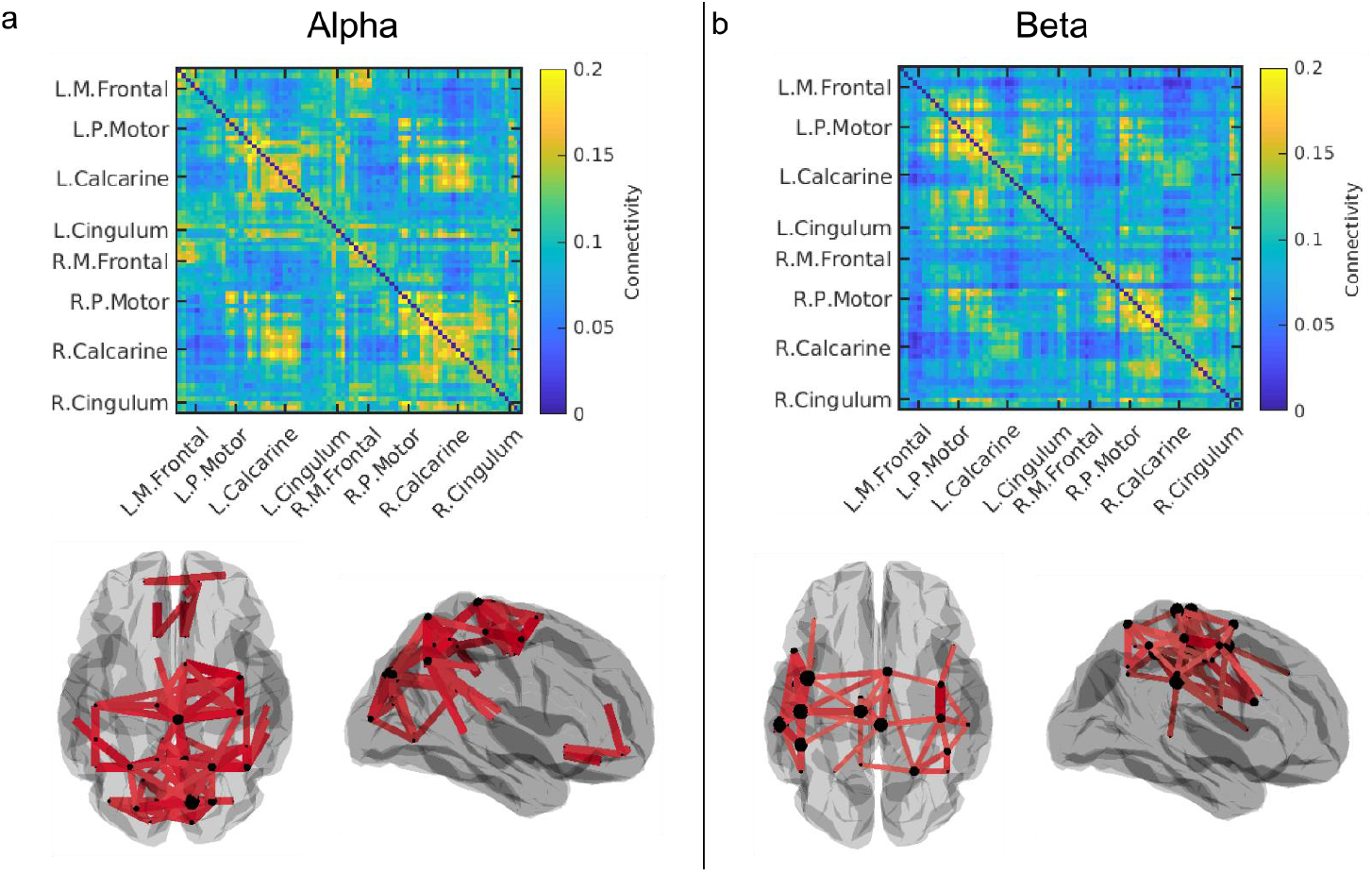
Resting-state connectivity plots derived from OPM data. Alpha- (a) and beta- (b) band connectivity matrices averaged across the 7 participants. Brain surfaces show the connections within 20 % (alpha) and 30 % (beta) of the maximum value.

Resting-state connectivity results, derived from cryogenic data, are plotted in Figure 6. Panels a and b show alpha- and beta-band connectivity matrices, respectively. In each panel, 9 different matrices are shown: these correspond to the 9 (randomly selected) groups of 7 subjects. Colour bars are the same for all matrices. The 3D brains show dominant connections (within 30 % of the maximum connectivity value). (These are derived from the grand average across 63 subjects.) Here we see that alpha oscillations mediate connections primarily in occipital areas whilst the beta-band connectome shows a more widespread connectivity, between occipital and parietal regions. Interestingly, whilst a common structure exists across all groups, in both bands, there is large discrepancy between connectivity strength values across groups.

**Figure 6:**
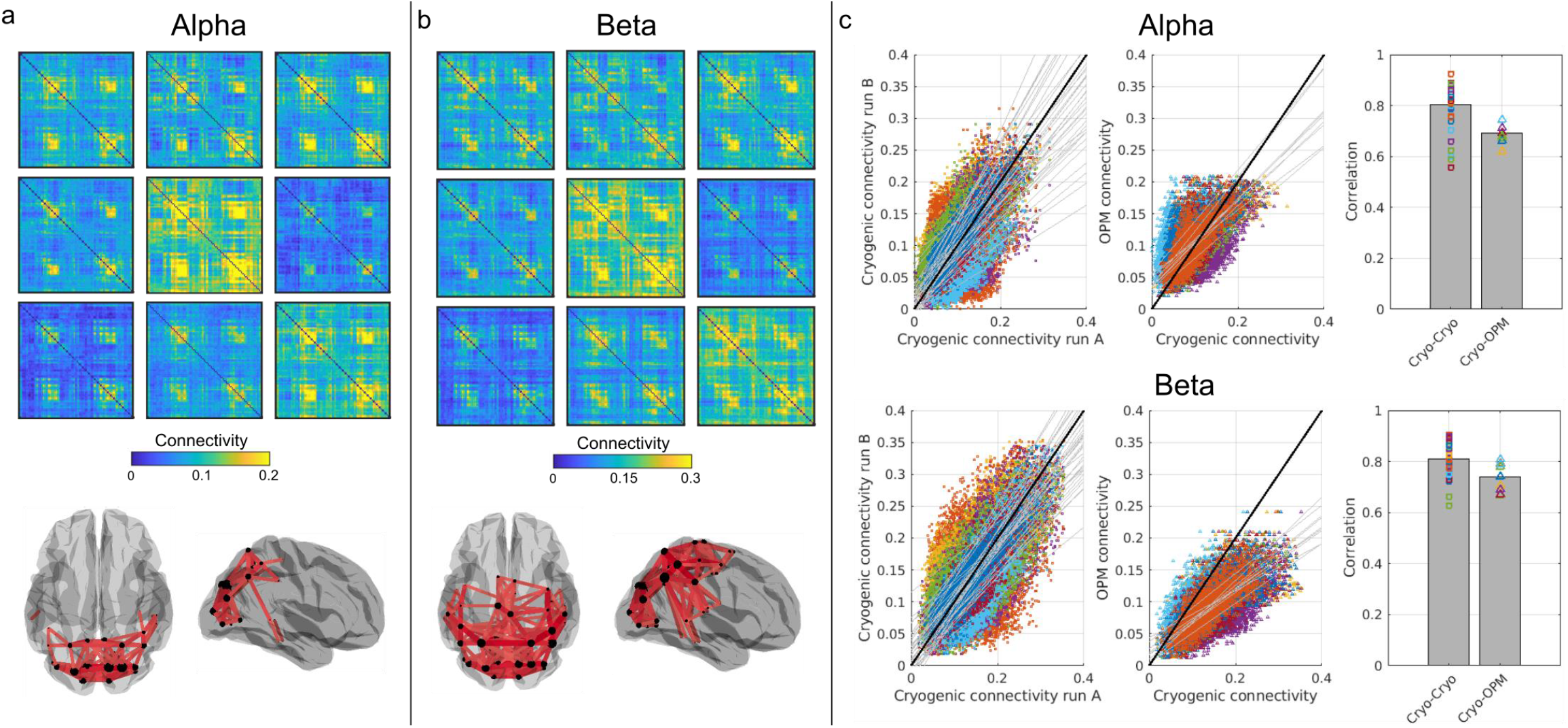
Resting-state group connectivity matrices from cryogenic data and a comparison with the OPM-derived connectome. Alpha- (a) and beta- (b) band connectivity matrices from 9 groups of 7 subjects. 3D brain plots show dominant connections (within 30 % of the maximum value). Note that even though these are group averaged results, clear difference across groups remain (although the overall pattern appears robust). c) The scatter plots on the left show cryogenic-derived connectivity values, with different groups plotted against each other i.e. each data point shows connectivity for the same connection, in two different subject groups, plotted against each other. The black line shows y=x; the coloured lines show lines of best fit for the 36 different possible comparisons between independent groups. The scatter plots in the centre show cryogenic-derived connectivity versus OPM-derived connectivity values. 9 separate comparisons are made between the OPM-derived connectome (averaged across 7 subjects) and 9 separate cryogenic-derived connectomes (each the average of 7 subjects). The bar plot shows mean correlation values for cryogenic-to-cryogenic connectivity (left-hand bar) and OPM-to-cryogenic connectivity (right-hand bar). Note that the scatter of points is relatively similar indicating good agreement between OPM- and cryogenic-derived resting-state connectivity.

Figure 6c shows a comparison between cryogenic- and OPM-derived resting-state connectomes. The upper and lower rows show results for alpha and beta bands, respectively. In the scatter plots, we show resting-state connectivity values from different groups of people, plotted against each other. In the left-hand scatter plot, 36 different comparisons are made, corresponding to 36 available pairs within our 9-group connectivity matrices. The black line shows ‘*y* = *x*’ and, given that the points represent connectivity values derived using the same system, for the same connections, in different subject groups, we would expect to see a scatter around this line. This is broadly the case, however it is important to note how wide the variation around this line is. This reflects differences between subjects. The middle scatter plot contains 9 comparisons. Here, our group-averaged OPM-derived connectivity values are plotted against equivalent values for each of the 9 cryogenic group averages. Note here that for both the alpha and beta band, although data do not necessarily lie along the ‘*y* = *x*’ line, a very clear linear trend is observed.

Finally, the right-hand bar chart shows correlation values between group-level connectomes; the left-hand bar shows cryogenic vs cryogenic connectomes; the right-hand bar shows OPM vs cryogenic connectomes. The bars show averages, whilst all data contributing to those averages are shown overlaid as squares/triangles. Here we see a clear result; if our OPM system was not able to successfully measure resting-state connectivity, we would expect the OPM vs cryogenic comparison to be significantly lower than the cryogenic vs cryogenic comparison. However, whilst the average is slightly lower, this effect is not significant, and overall it is clear that there is similarity between OPM-derived and cryogenic-derived resting-state functional connectivity measurements.

## 4 Discussion

OPMs represent a step change for MEG instrumentation: OPM-MEG offers the potential for cheaper MEG systems which can ultimately come into more widespread use, particularly in clinical settings. Wearable helmets mean that sensors move with the head, removing worries around subject movement which can lead to data becoming unusable in cryogenic systems (Boto et al., 2018). Flexible placement of small and lightweight sensors means that, in principle, an OPM-MEG system can adapt to any head shape or size (Hill et al., 2020). Ultimately this means that OPM-based systems are better able to accommodate challenging patient groups, in particular children (with smaller heads) or subject groups who find it hard to keep sufficiently still in a conventional scanning environment. The ability to move whilst scanning opens up new possibilities for neuroscientific experiment – for example we can scan people as they undertake naturalistic tasks (Hill et al., 2019) or become immersed in a virtual environment (Roberts et al., 2019). Finally, because OPM sensors can get closer to the brain, we can capture better data with higher sensitivity and spatial resolution (Boto et al., 2019, 2016; Iivanainen et al., 2017). These factors point towards OPMs superseding cryogenic MEG devices in the coming years. However, the technology remains largely unproven, and it is critical that OPM-MEG systems begin to demonstrate that they can perform at least as well as cryogenic systems for important neuroscientific measurements.

Functional connectivity is an area that has become of great importance in recent years. Canonical networks, and the functional connectivities that underlie them, are fundamental to healthy brain function and have been shown to be perturbed in a number of abnormalities ranging from mental health disorders that strike in the very young, to neurodegenerative conditions that become a problem for the elderly. The combination of high spatial and temporal resolution makes MEG, arguably, the technique of choice for measurement of brain network activity and connectivity. This is particularly true for the measurement of dynamic connectivity (e.g. during a task) where we might aim to probe the formation and dissolution of transient networks as they modulate to support cognition. It is for these reasons that functional connectivity and network measurement represent a key part of MEG research. Consequently, showing that OPM-MEG systems are capable of such measurements, with similar fidelity to conventional devices, is a key step forward in the journey towards a viable OPM-MEG device.

Here, we aimed to show that OPM-MEG could indeed offer complete characterisation of the brain-wide functional connectome. As noted in our introduction, such demonstration relies not only on high fidelity (unaveraged) MEG data but also on whole brain coverage and high spatial specificity; given the limited number of channels in OPM-MEG systems (∼50 compared to ∼300 in cryogenic systems) these latter points could have posed a challenge. However, results show that 50-channel OPM-MEG, in combination with accurate co-registration procedures and an appropriate source localisation algorithm, can indeed measure functional connectivity with similar efficacy to a cryogenic system.

Our task-based connectivity demonstration showed that both cryogenic- and OPM-MEG yield robust networks in the alpha, beta and gamma bands in response to a visuo-motor paradigm. Of interest here is the ability to measure individual differences between subjects. We showed that, in the beta band, both systems measured a robust sensori-motor network which exhibited significantly more bilateral connectivity in subject two, compared to subject 1. Of course, the reason for these differences between subjects is unclear, but the fact that the same (significant) differences were observed using both cryogenic and OPM-MEG is compelling. We argue that this finding is important for two reasons. First, from an OPM-MEG point of view, it validates the fact that the MEG data are of high quality; indeed the ability to “tell which subject you are scanning” based only on the MEG data is a satisfying demonstration of the equivalence between OPM and cryogenic derived data. Second, more broadly, this finding demonstrates the importance of inter-individual differences. It is rare in the MEG literature that single subjects are scanned multiple times; more usually, large subject groups containing tens or hundreds of subjects from different ‘groups’ are scanned and differences between those groups sought. Here, we show that sizeable differences between healthy individuals can be robustly observed in MEG data and it is tempting to speculate that these differences are larger than the more subtle deviations that are often observed between groups. It follows that future study should take this into account. Indeed, given the robustness of the within-subject measures that we observe here, it is likely that acquisition of longitudinal measures, tracking how e.g. a patient’s brain changes throughout the course of an illness, may ultimately be more fruitful (and more useful clinically) than cross sectional group studies.

In many ways, our resting-state data posed a greater challenge for OPM-MEG compared to task-based data, for the simple reason that the task-based connectome could be averaged over trials, potentially masking the effect of any artefacts at the sensor level. Conversely, resting state connectivity must be inferred based on unaveraged data, meaning that sensor artefacts could have a greater influence. Our findings showed that similar resting state network structure could be elucidated both using cryogenic and OPM-based MEG. Our beta band analyses showed that, when considering a group result across 7 subjects, cryogenic-derived connectome matrices showed 80% correlation; when comparing OPM and cryogenic derived connectomes, this was reduced marginally to 74%. In the alpha band, this reduction was somewhat larger with 80% correlation for cryogenic derived connectomes reducing to 68% for OPM-cryogenic comparison. These reductions are not surprising considering the vast differences between the systems – in particular the channel count. However, in both cases the reduction was not statistically significant. Indeed, when looking at the range of correlation values across different subject groups, these reductions were small and could easily be due to differences in the groups of participants scanned. These data, coupled with our task-based results, therefore show that OPM-MEG, even with a modest number of sensors, is able to effectively characterise the human connectome.

These resting state results are somewhat surprising: given the fact that OPMs are configured as magnetometers, as distinct from gradiometers, we might expect a higher degree of interference in our OPM compared to our cryogenic data. Consequently, one might expect a drop in fidelity of the connectivity assessment, particularly for the resting state where no averaging is possible. In fact, further analysis showed that magnetic artefacts of no interest are present in OPM-MEG sensor level data, however, using beamforming these artefacts are likely to be eliminated very efficiently. (See appendix, where the magnetic artefact of the heart has been close to be eliminated via beamforming.) This is an important point; it is hard to imagine how one might efficiently form an efficient gradiometer-based system from OPMs. Axial gradiometers would require OPMs to be stacked on top of one another, making the wearable helmet bulky and probably impractical. Planar gradiometers are possible to form and have been shown to be effective (Hill et al., 2019), however this involves a digital subtraction of signals from two adjacent OPMs which means (assuming a simple Gaussian model) a likely 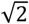 increase in sensor noise. With this in mind, it is positive that mechanisms such as beamforming work well. It remains to be seen as to whether other interference rejection strategies (for example signal source separation; (Taulu and Simola, 2006)) are also effective, but as shown in our appendix, effective interference minimisation will be extremely important for future OPM-MEG connectivity metrics.

Whilst OPM- and cryogenic-derived connectomes were largely similar, there are some differences which are worth noting. First, both in resting and task-positive connectivity, we noted generally weaker occipital connections for the OPM system. This is evident in Figure 2, where gamma-band results are noisier in the OPM system. It is also apparent in the resting-state measurements where the cryogenic system demonstrated higher occipital connectivity in alpha and beta bands. The reason for this difference is unclear, however we speculate that it results from the way in which the MEG helmet was fitted. The additively-manufactured helmet was a good fit for most subjects over the top of the head, but a noticeable gap (of ∼1– 2 cm) exists across the back of the head. This gap acts to move sensors away from occipital lobe and, consequently, signals will diminish in accordance with an inverse square law. In contrast, in a cryogenic MEG system most subjects tend to rest their head on the back of the scanner, meaning occipital lobe has the best coverage (and, correspondingly, frontal cortices the worst). It is therefore likely that a better fitting helmet (for example with sensor mounts that enable a degree of movement in the radial direction) would offer improved performance for OPM-based connectivity measures. Related to this, the alpha band resting state connectivity matrix showed higher connectivity in frontal regions for OPMs. Again this could be a proximity effect: a recent study by Coquelet et al., (2020) showed that electroencephalography (EEG) offered higher frontal cortex connectivity compared to (conventional) MEG and they cited an increased gap between the sensors and the brain in MEG, in frontal regions, as the reason. Our OPM system likely brings sensors closer to frontal cortices and this could explain the increased sensitivity to frontal alpha connectivity. What is clear is that inhomogeneous brain-to-sensor spacing for different areas of cortex can have a marked effect on results and this effect must be taken into account in future generations of scanner design. One potential solution is to use individual scanner-casts, however these tend to be difficult and time-consuming to generate and are expensive. The introduction of more sophisticated helmets that allow a degree of adaptation to different head shapes (e.g. by including built-in facility to adjust sensor positions along the radial direction), could negate this problem.

## 5 Conclusion

In conclusion, our study has shown that OPM-MEG can measure whole-brain functional connectivity with a fidelity similar to that demonstrated by conventional cryogenic MEG machines. In the resting state, our results show that connectome matrices from OPM and cryogenic systems exhibit an extremely high degree of similarity, with correlation values >70 %. This value is not measurably different to the correlation observed between connectomes measured across different subject groups on a single cryogenic MEG device. In a task-based study, we showed that robust differences in connectivity between individuals (scanned multiple times) exist, and similar individualised features could be identified in cryogenic and OPM-MEG measurements, again demonstrating the fidelity of OPM-MEG data. OPMs offer a step change for MEG instrumentation, however OPM-MEG remains a nascent technology with significant work still to be done. The present demonstration takes us one step closer to routine use of OPM-MEG for neuroscientific measurement. This adds weight to the argument that OPMs will ultimately supersede cryogenic-based instrumentation.

## Acknowledgements

We express our sincere thanks to collaborators at the Wellcome Centre for Human Neuroimaging, University College London, UK for extremely helpful discussions and ongoing support. Special thanks to Benjamin A.E. Hunt, who collected data from the 63 subjects as part of the UK MEG partnership programme, which we used here for comparison with OPM data. This work was supported by the UK Quantum Technology Hub in Sensing and Timing, funded by the Engineering and Physical Sciences Research Council (EPSRC) (EP/T001046/1), and a Wellcome Collaborative Award in Science (203257/Z/16/Z and 203257/B/16/Z) awarded to Gareth R. Barnes, RB and MJB.

## Appendix External interference and its influence on OPM data

A major concern related to connectivity measurement using OPM-MEG is the influence of external magnetic interference. As outlined in our introduction, most cryogenic MEG systems employ gradiometers to reduce the effect of external magnetic fields. Some also use reference arrays, higher-order synthetic gradiometry (Vrba and Robinson, 2001), or software approaches (Taulu and Simola, 2006). However, OPMs are inherently formed as magnetometers and this means that interference is more problematic. This is a particular concern for connectivity measures (in the resting state) since averaging across trials is impossible. Thus, understanding how OPM data is influenced by interference is important.

A good example of interference from a distal source is the magnetic artefact from the heart. Even at a distance, the field from the heart is many times larger than the field from the brain. Also, the fundamental frequencies of the heartbeat overlap with the frequency bands of interest for connectivity (i.e. the alpha and beta ranges). Consequently, this poses a significant challenge since, if the heartbeat artefact appears across many sensors, and is projected (via source localisation) into brain space, this will necessarily inflate functional connectivity measurement. For these reasons, we aimed to test the extent to which the heartbeat artefact is found in source-localised OPM-MEG data.

### Methods

In order to probe the presence of a heartbeat artefact in OPM-MEG resting-state data, independent component analysis (ICA) was applied to beta-band filtered sensor-level data. ICA was applied in the temporal dimension using the fastICA algorithm (Hyvärinen, 1999); we selected a sufficient number of independent components to explain 95 % of the data variance. This procedure was applied to data from all 7 subjects independently. For one subject, a clear heartbeat artefact was contained within a single independent component. For the other 6 subjects, the heartbeat was split across two independent components.

Following this, for all subjects we re-ran the beamformer spatial filter in order to reconstruct source-space data at the 78 pre-selected AAL regions. This was done in two different ways:

1. Following ICA, data were reconstructed with the heartbeat artefact removed (by removing either 1 or 2 independent components). The data covariance matrix was derived based on these reduced data, and a beamformer applied. Note that component removal in this way necessitates the application of matrix regularisation and so this was applied, using the Tikhonov method, with a regularisation parameter equal to 0.001 times the maximum singular value of the unregularised matrix.
2. Beamforming was applied without removal of the heartbeat artefact. This was done with no regularisation (as in our main manuscript) as well as with regularisation parameters equal to 5 % and 15 % of the maximum singular value of the unregularised matrix. (Note, the addition of regularisation reduces the ability of the beamformer to supress external magnetic interference, and consequently this proves a useful marker of how other source localisation algorithms (e.g. a dipole fit) might behave.)

In both cases we correlated the ICA-derived heartbeat to the source-localised MEG data in order to assess the influence of interference.

### Results and discussion

Figure A1a shows representative beta-band filtered OPM-MEG data at the channel level. Data for four channels, for a single subject, are shown and we see that the beta-band component of the heartbeat is easily identified. Figure A1 shows the cortical topography of correlation with the heartbeat artefact following source localisation. In the top left, we show the case where the heartbeat artefact was removed. The top right, bottom left, and bottom right show the case for a beamformer with zero, 5 % and 15 % regularisation respectively. As expected, heartbeat correlation is zero in the case where the heartbeat has been removed (a simple consequence of ICA). More importantly, correlation is very close to zero (0.03 ± 0.02 (mean ± standard deviation across all 78 regions)) when an unregularised beamformer is applied. As regularisation is increased, correlation increases markedly (to 0.06 ± 0.04 and 0.11 ± 0.06 for 5 % and 15 % regularisation, respectively). Note that correlation is most pronounced for deeper regions.

It is clear from this result that the good performance of OPM-MEG in connectivity assessment that we see in our manuscript is due, at least in part, to the noise rejection characteristics of the beamformer. In cases where beamforming is less efficacious, artefacts begin to leak into source-space data and as a consequence, connectivity measurement (or indeed any assessment of neural oscillatory processes) would become contaminated. Of course, the artefact as defined here can also be removed by other techniques (e.g. ICA – as demonstrated) but this relies on *a-priori* artefact identification. This was easy for the temporally well-characterised heartbeat, but is significantly harder for less well-known sources of magnetic interference. On the other hand, the beamformer does not rely on *a-priori* assumptions. Further, if it works well on the heartbeat, we can assume it works equally well in nulling other sources of external magnetic interference. In conclusion, we recommend that beamforming (or at least adaptive source-localisation techniques with good interference rejection properties) are used when attempting to assess source-space functional connectivity using OPM-MEG.

**Figure A1:**
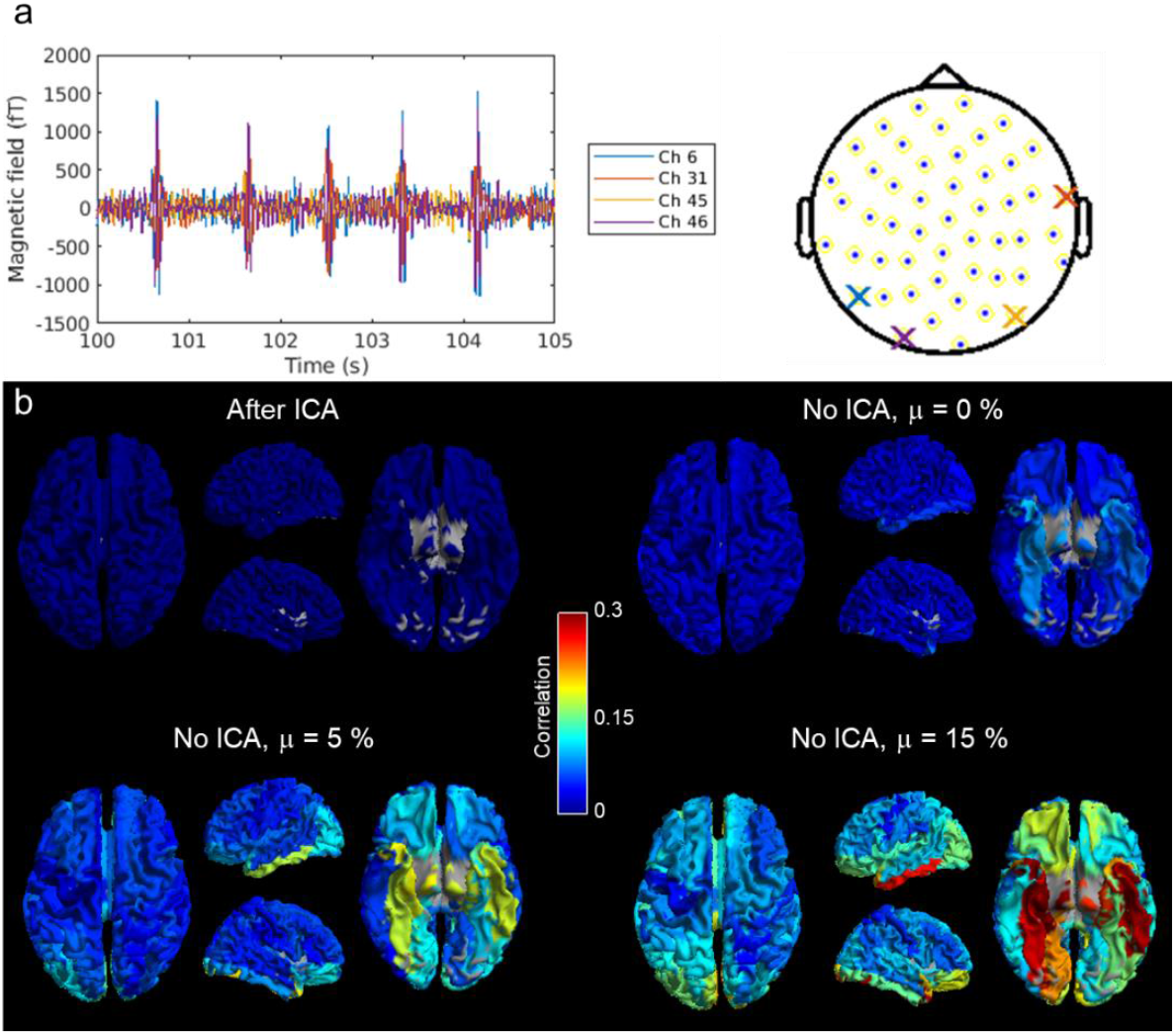
Presentation of the heartbeat artefact in OPM-MEG: a) Sensor-space data filtered to the beta band. Four channels are shown (indicated on the sensor layout on the right) and despite the separation of approximately 40 cm between the heart itself and the head-mounted sensors, the magnetocardiogram can be seen clearly. b) Correlation between the heartbeat artefact and source-localised data, across 78 AAL regions. Top left: Beamforming applied with the cardiac artefact removed. Top right: Beamforming applied to the full dataset with no regularisation. Bottom left: Beamforming applied with 5 % regularisation. Bottom right: Beamforming applied with 15 % regularisation.

## References

Altarev, I., Fierlinger, P., Lins, T., Marino, M.G., Nießen, B., Petzoldt, G., Reisner, M., Stuiber, S., Sturm, M., Taggart Singh, J., Taubenheim, B., Rohrer, H.K., Schläpfer, U., 2015. Minimizing magnetic fields for precision experiments. J. Appl. Phys. 117. https://doi.org/10.1063/1.4922671

Baker, A.P., Brookes, M.J., Rezek, I.A., Smith, S.M., Behrens, T., Smith, P.J.P., Woolrich, M., 2014. Fast transient networks in spontaneous human brain activity. Elife 2014, e01867. https://doi.org/10.7554/eLife.01867

Barry, D.N., Tierney, T.M., Holmes, N., Boto, E., Roberts, G., Leggett, J., Bowtell, R., Brookes, M.J., Barnes, G.R., Maguire, E.A., 2019. Imaging the human hippocampus with optically-pumped magnetoencephalography. Neuroimage 203. https://doi.org/10.1016/j.neuroimage.2019.116192

Beckmann, C.F., DeLuca, M., Devlin, J.T., Smith, S.M., 2005. Investigations into resting-state connectivity using independent component analysis. Philos. Trans. R. Soc. B Biol. Sci. 360, 1001–1013. https://doi.org/10.1098/rstb.2005.1634

Benjamini, Y., Hochberg, Y., 1995. Controlling the False Discovery Rate: A Practical and Powerful Approach to Multiple Testing. J. R. Stat. Soc. Ser. B. https://doi.org/10.1111/j.2517-6161.1995.tb02031.x

Biswal, B., Zerrin Yetkin, F., Haughton, V.M., Hyde, J.S., 1995. Functional connectivity in the motor cortex of resting human brain using echo‐planar mri. Magn. Reson. Med. 34, 537–541. https://doi.org/10.1002/mrm.1910340409

Borna, A., Carter, T.R., Colombo, A.P., Jau, Y.Y., McKay, J., Weisend, M., Taulu, S., Stephen, J.M., Schwindt, P.D.D., 2020. Non-invasive functional-brain-imaging with an OPM-based magnetoencephalography system. PLoS One 15. https://doi.org/10.1371/journal.pone.0227684

Boto, E., Bowtell, R., Krüger, P., Fromhold, T.M., Morris, P.G., Meyer, S.S., Barnes, G.R., Brookes, M.J., 2016. On the potential of a new generation of magnetometers for MEG: A beamformer simulation study. PLoS One 11, e0157655. https://doi.org/10.1371/journal.pone.0157655

Boto, E., Holmes, N., Leggett, J., Roberts, G., Shah, V., Meyer, S.S., Muñoz, L.D., Mullinger, K.J., Tierney, T.M., Bestmann, S., Barnes, G.R., Bowtell, R., Brookes, M.J., 2018. Moving magnetoencephalography towards real-world applications with a wearable system. Nature 555, 657–661. https://doi.org/10.1038/nature26147

Boto, E., Holmes, N., Tierney, T.M., Leggett, J., Hill, R., Mellor, S., Roberts, G., Barnes, G.R., Bowtell, R., Brookes, M.J., Boto, E., Holmes, N., Tierney, T.M., Leggett, J., Hill, R., Mellor, S., Roberts, G., Barnes, G.R., Bowtell, R., Brookes, M.J., 2020. Magnetoencephalography Using Optically Pumped Magnetometers, in: Fifty Years of Magnetoencephalography. Oxford University Press, New York, pp. 104–124. https://doi.org/10.1093/oso/9780190935689.003.0008

Boto, E., Meyer, S.S., Shah, V., Alem, O., Knappe, S., Kruger, P., Fromhold, T.M., Lim, M., Glover, P.M., Morris, P.G., Bowtell, R., Barnes, G.R., Brookes, M.J., 2017. A new generation of magnetoencephalography: Room temperature measurements using optically-pumped magnetometers. Neuroimage 149, 404–414. https://doi.org/10.1016/j.neuroimage.2017.01.034

Boto, E., Seedat, Z.A., Holmes, N., Leggett, J., Hill, R.M., Roberts, G., Shah, V., Fromhold, T.M., Mullinger, K.J., Tierney, T.M., Barnes, G.R., Bowtell, R., Brookes, M.J., 2019. Wearable neuroimaging: Combining and contrasting magnetoencephalography and electroencephalography. Neuroimage 201. https://doi.org/10.1016/j.neuroimage.2019.116099

Bright, M.G., Whittaker, J.R., Driver, I.D., Murphy, K., 2020. Vascular physiology drives functional brain networks. Neuroimage 217. https://doi.org/10.1016/j.neuroimage.2020.116907

Brookes, M.J., Vrba, J., Robinson, S.E., Stevenson, C.M., Peters, A.M., Barnes, G.R., Hillebrand, A., Morris, P.G., 2008. Optimising experimental design for MEG beamformer imaging. Neuroimage 39, 1788–1802. https://doi.org/10.1016/j.neuroimage.2007.09.050

Brookes, M.J., Woolrich, M.W., Barnes, G.R., 2012. Measuring functional connectivity in MEG: A multivariate approach insensitive to linear source leakage. Neuroimage 63, 910–920. https://doi.org/10.1016/j.neuroimage.2012.03.048

Brookes, M.J., Woolrich, M.W., Luckhoo, H., Price, D., Hale, J.R., Stephenson, M.C., Barnes, G.R., Smith, S.M., Morris, P.G., 2011. Investigating the electrophysiological basis of resting state networks using magnetoencephalography. Proc. Natl. Acad. Sci. 108, 16783–16788. https://doi.org/10.1073/pnas.1112685108

Cohen, D., 1972. Magnetoencephalography: Detection of the brain’s electrical activity with a superconducting magnetometer. Science (80-.). 175, 664–666. https://doi.org/10.1126/science.175.4022.664

Coquelet, N., De Tiège, X., Destoky, F., Roshchupkina, L., Bourguignon, M., Goldman, S., Peigneux, P., Wens, V., 2020. Comparing MEG and high-density EEG for intrinsic functional connectivity mapping. Neuroimage 210. https://doi.org/10.1016/j.neuroimage.2020.116556

Engel, A.K., Gerloff, C., Hilgetag, C.C., Nolte, G., 2013. Intrinsic Coupling Modes: Multiscale Interactions in Ongoing Brain Activity. Neuron 80, 867–886. https://doi.org/10.1016/j.neuron.2013.09.038

Fox, M.D., Raichle, M.E., 2007. Spontaneous fluctuations in brain activity observed with functional magnetic resonance imaging. Nat. Rev. Neurosci. https://doi.org/10.1038/nrn2201

Gross, J., Kujala, J., Hämäläinen, M.S., Timmermann, L., Schnitzler, A., Salmelin, R., 2001. Dynamic imaging of coherent sources: Studying neural interactions in the human brain. Proc. Natl. Acad. Sci. 98, 694–699. https://doi.org/10.1073/pnas.98.2.694

Hill, R.M., Boto, E., Holmes, N., Hartley, C., Seedat, Z.A., Leggett, J., Roberts, G., Shah, V., Tierney, T.M., Woolrich, M.W., Stagg, C.J., Barnes, G.R., Bowtell, R.R., Slater, R., Brookes, M.J., 2019. A tool for functional brain imaging with lifespan compliance. Nat. Commun. 10. https://doi.org/10.1038/s41467-019-12486-x

Hill, R.M., Boto, E., Rea, M., Holmes, N., Leggett, J., Coles, L.A., Papastavrou, M., Everton, S.K., Hunt, B.A.E., Sims, D., Osborne, J., Shah, V., Bowtell, R., Brookes, M.J., 2020. Multi-channel whole-head OPM-MEG: Helmet design and a comparison with a conventional system. Neuroimage. https://doi.org/10.1016/j.neuroimage.2020.116995

Hipp, J.F., Hawellek, D.J., Corbetta, M., Siegel, M., Engel, A.K., 2012. Large-scale cortical correlation structure of spontaneous oscillatory activity. Nat. Neurosci. 15, 884–890. https://doi.org/10.1038/nn.3101

Holmes, N., Leggett, J., Boto, E., Roberts, G., Hill, R.M., Tierney, T.M., Shah, V., Barnes, G.R., Brookes, M.J., Bowtell, R., 2018. A bi-planar coil system for nulling background magnetic fields in scalp mounted magnetoencephalography. Neuroimage 181, 760–774. https://doi.org/10.1016/j.neuroimage.2018.07.028

Homölle, S., Oostenveld, R., 2019. Using a structured-light 3D scanner to improve EEG source modeling with more accurate electrode positions. J. Neurosci. Methods 326, 108378. https://doi.org/10.1016/j.jneumeth.2019.108378

Hoogenboom, N., Schoffelen, J.M., Oostenveld, R., Parkes, L.M., Fries, P., 2006. Localizing human visual gamma-band activity in frequency, time and space. Neuroimage. https://doi.org/10.1016/j.neuroimage.2005.08.043

Hunt, B.A.E., Tewarie, P.K., Mougin, O.E., Geades, N., Jones, D.K., Singh, K.D., Morris, P.G., Gowland, P.A., Brookes, M.J., 2016. Relationships between cortical myeloarchitecture and electrophysiological networks. Proc. Natl. Acad. Sci. 113, 13510–13515. https://doi.org/10.1073/pnas.1608587113

Hutchison, R.M., Womelsdorf, T., Allen, E.A., Bandettini, P.A., Calhoun, V.D., Corbetta, M., Della Penna, S., Duyn, J.H., Glover, G.H., Gonzalez-Castillo, J., Handwerker, D.A., Keilholz, S., Kiviniemi, V., Leopold, D.A., de Pasquale, F., Sporns, O., Walter, M., Chang, C., 2013. Dynamic functional connectivity: Promise, issues, and interpretations. Neuroimage 80, 360–378. https://doi.org/10.1016/j.neuroimage.2013.05.079

Hyvärinen, A., 1999. Fast and robust fixed-point algorithms for independent component analysis. IEEE Trans. Neural Networks 10, 626–634. https://doi.org/10.1109/72.761722

Iivanainen, J., Stenroos, M., Parkkonen, L., 2017. Measuring MEG closer to the brain: Performance of on-scalp sensor arrays. Neuroimage 147, 542–553. https://doi.org/10.1016/j.neuroimage.2016.12.048

Iivanainen, J., Zetter, R., Parkkonen, L., 2019. Potential of on-scalp MEG: Robust detection of human visual gamma-band responses. bioRxiv 602342. https://doi.org/10.1101/602342

Johnson, C.N., Schwindt, P.D.D., Weisend, M., 2013. Multi-sensor magnetoencephalography with atomic magnetometers. Phys. Med. Biol. 58, 6065–6077. https://doi.org/10.1088/0031-9155/58/17/6065

Kamada, K., Sato, D., Ito, Y., Natsukawa, H., Okano, K., Mizutani, N., Kobayashi, T., 2015. Human magnetoencephalogram measurements using newly developed compact module of high-sensitivity atomic magnetometer. Jpn. J. Appl. Phys. 54, 026601. https://doi.org/10.7567/JJAP.54.026601

Kim, K., Begus, S., Xia, H., Lee, S.K., Jazbinsek, V., Trontelj, Z., Romalis, M. V., 2014. Multi-channel atomic magnetometer for magnetoencephalography: A configuration study. Neuroimage 89, 143–151. https://doi.org/10.1016/j.neuroimage.2013.10.040

Luckhoo, H., Hale, J.R., Stokes, M.G., Nobre, A.C., Morris, P.G., Brookes, M.J., Woolrich, M.W., 2012. Inferring task-related networks using independent component analysis in magnetoencephalography. Neuroimage 62, 530–541. https://doi.org/10.1016/j.neuroimage.2012.04.046

Menon, V., 2011. Large-scale brain networks and psychopathology: A unifying triple network model. Trends Cogn. Sci. https://doi.org/10.1016/j.tics.2011.08.003

O’Neill, G.C., Bauer, M., Woolrich, M.W., Morris, P.G., Barnes, G.R., Brookes, M.J., 2015. Dynamic recruitment of resting state sub-networks. Neuroimage 115, 85–95. https://doi.org/10.1016/j.neuroimage.2015.04.030

Osborne, J., Orton, J., Alem, O., Shah, V., 2018. Fully integrated, standalone zero field optically pumped magnetometer for biomagnetism. Steep Dispers. Eng. Opto-Atomic Precis. Metrol. XI 10548.

Raichle, M.E., 2009. A paradigm shift in functional brain imaging. J. Neurosci. https://doi.org/10.1523/JNEUROSCI.4366-09.2009

Roberts, G., Holmes, N., Alexander, N., Boto, E., Leggett, J., Hill, R.M., Shah, V., Rea, M., Vaughan, R., Maguire, E.A., Kessler, K., Beebe, S., Fromhold, M., Barnes, G.R., Bowtell, R., Brookes, M.J., 2019. Towards OPM-MEG in a virtual reality environment. Neuroimage 199, 408–417. https://doi.org/10.1016/j.neuroimage.2019.06.010

Sander, T.H., Preusser, J., Mhaskar, R., Kitching, J., Trahms, L., Knappe, S., 2012. Magnetoencephalography with a chip-scale atomic magnetometer. Biomed. Opt. Express 3, 981. https://doi.org/10.1364/BOE.3.000981

Seedat, Z.A., Quinn, A.J., Vidaurre, D., Liuzzi, L., Gascoyne, L.E., Hunt, B.A.E., O’Neill, G.C., Pakenham, D.O., Mullinger, K.J., Morris, P.G., Woolrich, M.W., Brookes, M.J., 2020. The role of transient spectral ‘bursts’ in functional connectivity: A magnetoencephalography study. Neuroimage 209. https://doi.org/10.1016/j.neuroimage.2020.116537

Smith, S.M., Fox, P.T., Miller, K.L., Glahn, D.C., Fox, P.M., Mackay, C.E., Filippini, N., Watkins, K.E., Toro, R., Laird, A.R., Beckmann, C.F., 2009. Correspondence of the brain’s functional architecture during activation and rest. Proc. Natl. Acad. Sci. U. S. A. 106, 13040–13045. https://doi.org/10.1073/pnas.0905267106

Taulu, S., Simola, J., 2006. Spatiotemporal signal space separation method for rejecting nearby interference in MEG measurements. Phys. Med. Biol. 51, 1759–1768. https://doi.org/10.1088/0031-9155/51/7/008

Tierney, T.M., Holmes, N., Mellor, S., López, J.D., Roberts, G., Hill, R.M., Boto, E., Leggett, J., Shah, V., Brookes, M.J., Bowtell, R., Barnes, G.R., 2019. Optically pumped magnetometers: From quantum origins to multi-channel magnetoencephalography. Neuroimage In Press. https://doi.org/10.1016/j.neuroimage.2019.05.063

Tierney, T.M., Holmes, N., Meyer, S.S., Boto, E., Roberts, G., Leggett, J., Buck, S., Duque-Muñoz, L., Litvak, V., Bestmann, S., Baldeweg, T., Bowtell, R., Brookes, M.J., Barnes, G.R., 2018. Cognitive neuroscience using wearable magnetometer arrays: Non-invasive assessment of language function. Neuroimage 181, 513–520. https://doi.org/10.1016/j.neuroimage.2018.07.035

Tzourio-Mazoyer, N., Landeau, B., Papathanassiou, D., Crivello, F., Etard, O., Delcroix, N., Mazoyer, B., Joliot, M., 2002. Automated anatomical labeling of activations in SPM using a macroscopic anatomical parcellation of the MNI MRI single-subject brain. Neuroimage 15, 273–289. https://doi.org/10.1006/nimg.2001.0978

Vrba, J., Robinson, S.E., 2001. Signal processing in magnetoencephalography. Methods 25, 249–271. https://doi.org/10.1006/meth.2001.1238

Xia, H., Ben-Amar Baranga, A., Hoffman, D., Romalis, M. V., 2006. Magnetoencephalography with an atomic magnetometer. Appl. Phys. Lett. 89, Article number 211104. https://doi.org/10.1063/1.2392722

Zetter, R., Iivanainen, J., Parkkonen, L., 2019. Optical Co-registration of MRI and On-scalp MEG. Sci. Rep. 9, 5490. https://doi.org/10.1038/s41598-019-41763-4

